# On-grid labeling method for freeze-fracture replicas

**DOI:** 10.1101/2022.08.16.504048

**Authors:** Hiroko Osakada, Toyoshi Fujimoto

**Author notes:** Corresponding author: Toyoshi Fujimoto.

## Abstract

Sodium dodecyl sulfate–treated freeze-fracture replica labeling (SDS-FRL) is an electron microscopic (EM) method that can define the two-dimensional distribution of membrane proteins and lipids in a quantitative manner. Despite its unsurpassed merit, SDS-FRL has been adopted in a limited number of labs, probably because it requires a laborious labeling process as well as equipment and technique for freeze-fracture. Here, we present a method that reduces the manual labor significantly by mounting freeze-fracture replicas on EM grids prior to labeling. This was made possible by the discovery that freeze-fracture replicas invariably adhere to the carbon-coated formvar membrane with their platinum-carbon side, ensuring that the membrane molecules retained in replicas are accessible to labeling solutions. The replicas mounted on EM grids can be stored dry until labeling, checked by light microscopy before labeling, and labeled in the same manner as tissue sections. This on-grid method will make SDS-FRL easier to access for many researchers.

---

SDS-FRL is an electron microscopic method that employs freeze-fracture replicas and can define the two-dimensional distribution of membrane proteins and lipids in a quantitative manner [1,2]. This method can also analyze intracellular membranes by distinguishing their two leaflets and, when combined with quick-freezing, chemical fixation that may induce artificial molecular redistribution can be avoided. These advantages of SDS-FRL are well recognized and have been utilized to define the nanoscale distribution of both proteins and lipids [3-5]. Nevertheless, the number of labs using this method has been relatively small.

Two major obstacles are thought to deter labs from adopting the SDS-FRL for their work. The first is the need for the equipment and technique for freeze-fracture replica preparation, and the second is the laborious process of replica labeling, which appears difficult to perform simply by referring to manuals. With regard to the first obstacle, although some training is necessary for freeze-fracture, recent equipment is more user-friendly than it used to be. Additionally, as discussed later, the method described here will make it easier to collaborate with labs already familiar with freeze-fracture.

The second obstacle is more problematic than it may sound. This is because replicas treated with SDS tend to be fragmented into small pieces, often less than 100 µm in size, and need to be transferred between solutions multiple times under a stereomicroscope (Fig. 1A) [6]. At the end of labeling, replicas need to be picked up onto EM grids for observation, but this is the most time-consuming step and, even with skilled hands, some replicas may be lost as a result of sticking to grid edges, folding, or other reasons. Moreover, because the fracturing of frozen specimens occurs in random planes, replicas may turn out to contain no cellular structure but only fractured ice. A combination of these problems makes SDS-FRL appear daunting to the novice.

**Fig. 1.**
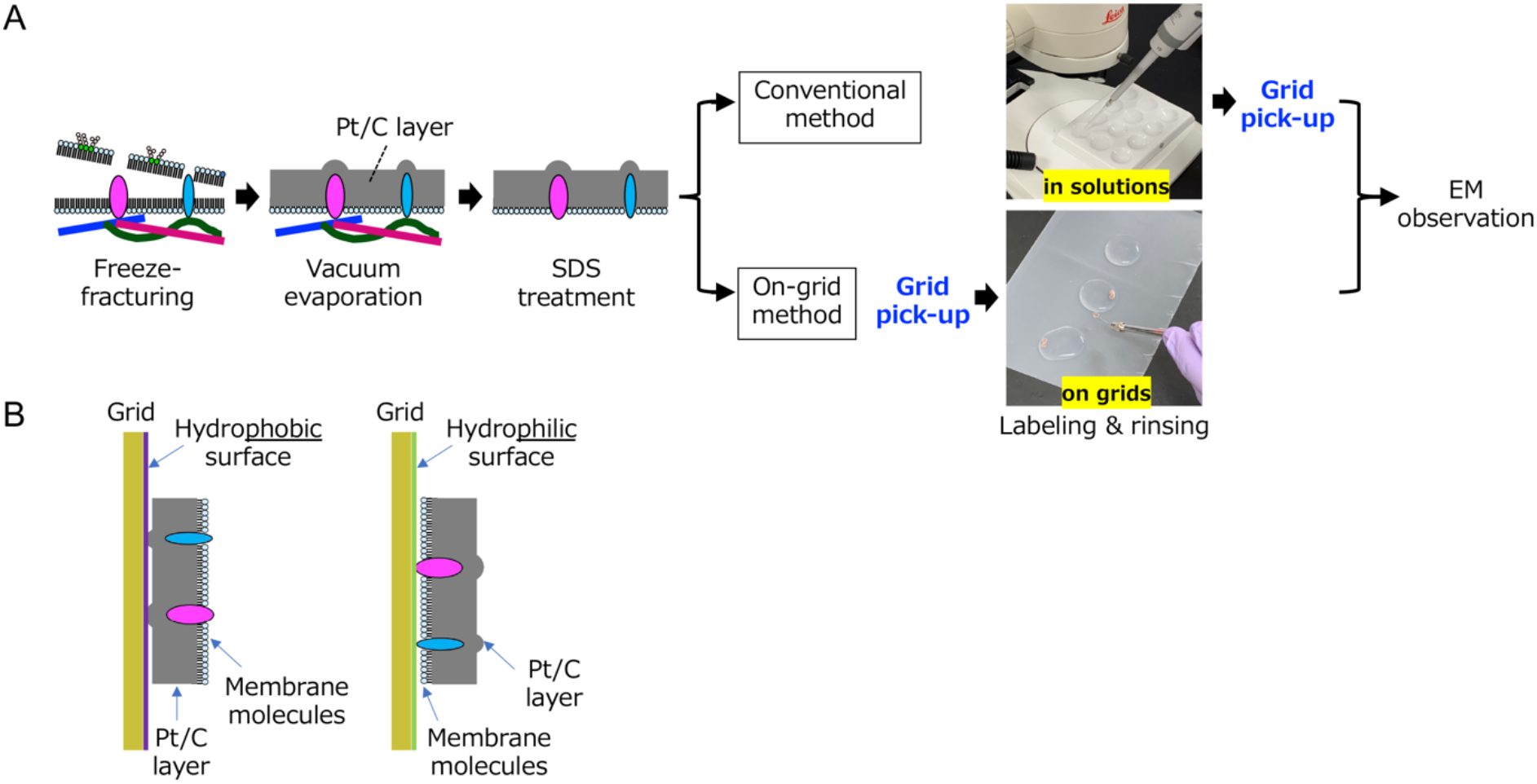
The on-grid SDS-FRL method. A. Comparison of the conventional method and the on-grid method. In the conventional method, freeze-fracture replicas are handled in solutions and picked up on EM grids in the end, whereas in the on-grid method, replicas are picked up on EM grids first and then labeled. B. The contrasting sidedness of freeze-fracture replicas picked up on the hydrophobic (carbon-coated) and hydrophilic (plasma-bombarded) surfaces.

Many of the above problems can be avoided or eased significantly if replicas are picked up on EM grids first and labeled thereafter. To enable this, replicas need to be mounted to EM grids facing their platinum–carbon side to the grid, so that their membrane molecule side is accessible to labeling solutions. The two sides of the replicas cannot be distinguished by light microscopy. A method of gluing replicas to grids with a chemical adhesive before SDS treatment has been reported, but this needs skill and subsequent removal of the adhesive [7]. Here, we presumed that the side of replicas adhering to grids may depend on the surface property of the latter; specifically, the hydrophobic platinum–carbon side and the hydrophilic membrane molecule side of replicas may bind preferentially to the hydrophobic and the hydrophilic surface, respectively (Fig. 1B).

To test this idea, we prepared formvar membrane on Maxtaform Copper/Rhodium grids (M100-CR, Electron Microscopy Sciences) and treated them in the following two different ways. 1) Carbon coating to make the surface hydrophobic. The grids were placed in CADE-4T (Meiwafosis) and treated for a few seconds to execute a thin carbon coating until the underlying filter paper became light gray; 2) Plasma ion bombardment to make the surface hydrophilic. The grids were treated for 2 s at an electric current of approximately 3 mA in PIB-10 (Vacuum Device). Freeze-fracture replicas of quick-frozen budding yeast (*Saccharomyces cerevisiae*, SEY6210 strain) that were prepared as described before [8] were used as test material. After overnight digestion in 2.5% SDS in phosphate-buffered saline (PBS) at 60ºC, the replicas were rinsed five times with 0.1% Tween 20 in PBS, rinsed twice in distilled water, and picked up on the grids treated in the two different ways as above. After drying to make the replicas firmly adhere to the formvar membrane, the grids were placed for 30 min on a drop of a blocking solution with the replica-mounted side facing downward. They were then transferred to subsequent solutions using a Nichrome wire loop attached to a holder (Fig. 1A). Labeling of a phospholipid, phosphatidylinositol 3-phosphate (PtdIns3P), was performed by successive incubation with recombinant glutathione-S-transferase (GST)-tagged p40^phox^ phox-homology (PX) domain, rabbit anti-GST antibody (A190-122A, Bethyl), and protein A-gold (10 nm) (PAG10, The University Medical Center Utrecht) [8]. For detection of phosphatidylcholine (PC) that was metabolically labeled with propargylcholine, biotin-azide (baseclick) was reacted with the alkyne residue in the PC headgroup by click chemistry, and the conjugated biotin was immunolabeled with mouse anti-biotin antibody (200-002-211, Jackson ImmunoResearch) and PAG10 [9]. To test protein labeling, Erg1-GFP expressed under the endogenous promoter was labeled by rabbit anti-GFP antibody (a gift from Dr. Masahiko Watanabe) and PAG10. The labeled replicas were observed under a transmission electron microscope (JEM1400EX, JEOL).

We first tested the sidedness of grid-mounted replicas using PtdIns3P labeling and found that all of the replicas picked up on the carbon-coated hydrophobic grids were labeled positively whereas those picked up on the plasma ion-bombarded hydrophilic grids were not. This result showed that replicas adhere to the hydrophobic surface with their platinum-carbon side. It was also notable that, even though the replicas picked up on the grids were completely dried before labeling, PtdIns3P was labeled specifically in the cytoplasmic leaflet of the vacuolar membrane, as we reported previously (Fig. 2A) [8]. The intensity of labeling was comparable to that in the replicas labeled by the conventional method.

**Fig. 2.**
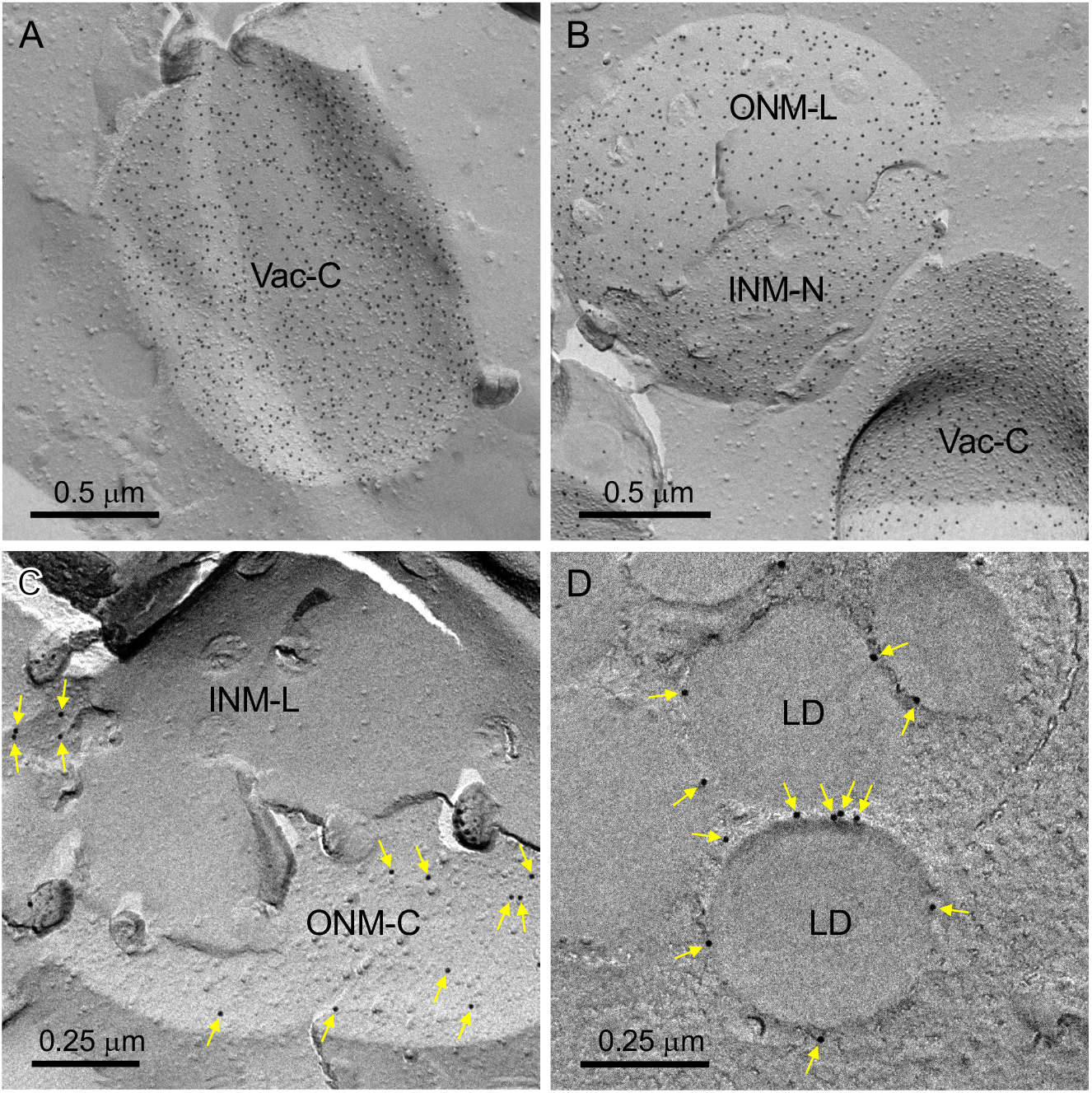
Representative results of the on-grid method. A. PtdIns3P in the cytoplasmic leaflet of the vacuole (Vac-C). B. Metabolically labeled PC in the cytoplasmic leaflet of the vacuole (Vac-C), the luminal leaflet of the outer nuclear membrane (ONM-L), and the nucleoplasmic leaflet of the inner nuclear membrane (INM-N). C. Erg1-GFP (yellow arrows) in the cytoplasmic leaflet of the outer nuclear membrane (ONM-C). ONM-L, the luminal leaflet of the inner nuclear membrane. D. Erg1-GFP (yellow arrows) in lipid droplets (LD). Note that LDs are cross-fractured.

PC labeling also took place successfully in replicas mounted on EM grids (Fig. 2B) [9]. The labeling specificity was confirmed by the elimination of labeling when copper sulfate was omitted from the click reaction solution (data not shown). The result indicates that labeling using click chemistry can also be performed by the on-grid method.

Next, the immunolabeling of proteins by the on-grid method was tested using yeast expressing Erg1-GFP. Erg1 is an integral membrane protein with both the amino- and carboxyl-termini facing the cytoplasm and distributes in the nuclear membrane, the endoplasmic leaflet, and lipid droplets [10]. Consistently, labels with anti-GFP antibody were observed in the cytoplasmic leaflet of the membranes and around lipid droplets (Figs. 2C, D). The result obtained by the conventional method was essentially the same (data not shown).

The result showed that replicas mounted to carbon-coated EM grids are labeled for PtdIns3P, metabolically labeled PC, and Erg1-GFP in the same manner as those labeled in the conventional manner. We confirmed that the grid-mounted replicas stored in a refrigerator for more than a week are labeled similarly. It is not surprising that dried replicas can be labeled, because membrane sheets (e.g., nitrocellulose, PVDF) blotted with lipids and proteins are used after drying for assays like protein-lipid overlay and Western blotting. Nevertheless, replicas differ from membrane sheets in that lipids and proteins are retained in a halved membrane structure so that whether the on-grid method works for other lipids and proteins needs to be tested individually.

In addition to the ease of handling, the grid-mounted replicas are advantageous in that they can be examined under a conventional upright light microscope to determine whether they contain cellular structures. For example, yeast cell pellets are observed as a cluster of round structures and can be distinguished from replicas that do not contain cells (Fig. 3). By discarding the latter replicas before labeling, time, labor, and reagents can be saved.

**Fig. 3.**
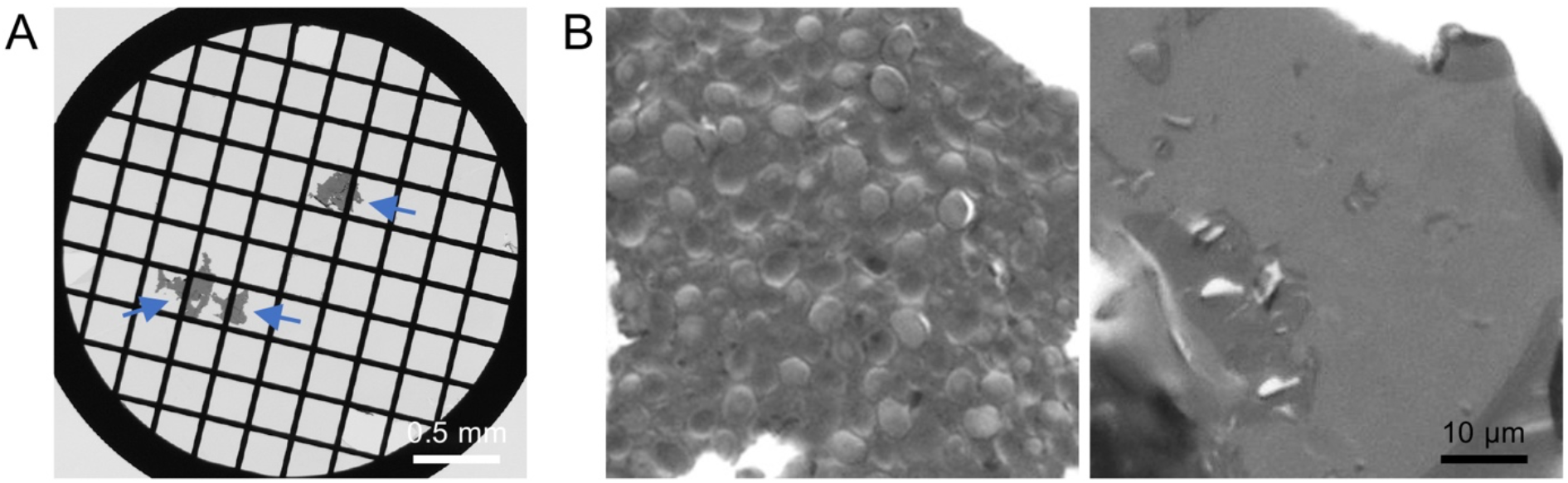
Grid-mounted freeze-fracture replicas by conventional light microscopy. A. A formvar membrane–coated EM grid (100 mesh) mounted with freeze-fracture replicas (blue arrows). B. Freeze-fracture replicas containing yeast cell pellet (left) and no cellular material (right).

Because the labeling procedure for on-grid replicas is essentially the same as that for sections, it can be adopted readily by labs familiar with conventional EM methods. These labs may be able to obtain grid-mounted replicas from collaborators who can prepare freeze-fracture replicas. For example, grid-mounted replicas of liposomes in different molecular compositions can be used to examine whether a lipid-binding protein binds to its potential target. We hope that the method reported here will make SDS-FRL accessible for more researchers.

## Acknowledgments

We thank Dr. Masahiko Watanabe (Hokkaido University) for the anti-GFP antibody gift. This study was supported by a Grant-in-Aid for Scientific Research from the Japan Society of the Promotion of Science (22H00446), by the Project for Elucidating and Controlling Mechanisms of Aging and Longevity from the Japan Agency for Medical Research and Development (21gm5010003), and by grants from the Nakatani Foundation for Advancement of Measuring Technologies in Biomedical Engineering and the Takeda Science Foundation to T.F.

